# Data-driven modeling of cholinergic modulation of neural microcircuits: bridging neurons, synapses, and network states

**DOI:** 10.1101/323865

**Authors:** Srikanth Ramaswamy, Henry Markram

**Affiliations:** Blue Brain Project, EPFL Biotech Campus, Geneva, Switzerland

**Keywords:** **Neuromodulation**, **acetylcholine**, **neocortex**, **microcircuits**, **cellular excitability**, **synaptic transmission**, **network states**

## Abstract

Neuromodulators, such as acetylcholine (ACh), control information processing in neural microcircuits by regulating neuronal and synaptic physiology. Computational models and simulations enable predictions on the potential role of ACh in reconfiguring network states. As a prelude into investigating how the cellular and synaptic effects of ACh collectively influence emergent network dynamics, we developed a data-driven framework incorporating phenomenological models of the anatomy and physiology of cholinergic modulation of the neocortex. The first-draft models were integrated into a biologically detailed tissue model of neocortical microcircuitry to predict how ACh affects different types of neurons and synapses, and consequently alters global network states. Preliminary simulations not only corroborate the long-standing notion that ACh desynchronizes network activity, but also reveal a potentially finegrained control over a spectrum of neocortical states. We show that low levels of ACh, such as those during sleep, drive microcircuit activity into slow oscillations and network synchrony, whereas high ACh concentrations, such as those during wakefulness, govern fast oscillations and network asynchrony. In addition, network states modulated by ACh levels shape spike-time cross-correlations across distinct neuronal populations in strikingly different ways. These effects are likely due to the differential regulation of neurons and synapses caused by increasing levels of ACh that enhances cellular excitability by increasing neuronal activity and decreases the efficacy of local synaptic transmission by altering neurotransmitter release probability. We conclude by discussing future directions to refine the biological accuracy of the framework, which will extend its utility and foster the development of hypotheses to investigate the role of neuromodulation in neural information processing.

## Introduction

Neuromodulatory axons projecting from the nucleus basalis of Meynert (NBM) in the basal forebrain densely innervate the neocortex that act through the release of ACh (Gielow and Zaborszky, 2017; Levey et al., 1987; Mesulam et al., 1983). Diffused release of ACh targets neurons and synapses in neocortical microcircuits, and regulates behavioral states, such as attention, wakefulness, learning, and memory (Hasselmo, 1995, 1999; Lee and Dan, 2012; Metherate et al., 1992). It is thought that the actions of ACh on the physiology of neurons and synapses plays a key role in switching cortical rhythms that underlie a diversity of behavioral states (Juliano and Jacobs, 1995; McCormick, 1992; Picciotto et al., 2012; Steriade et al., 1993; Xiang et al., 1998; Zagha and McCormick, 2014).

Much of our knowledge on the regulation of neuronal and synaptic physiology by ACh comes from studies in cortical slices that have combined whole-cell somatic recordings and iontophoretic application of ACh agonists, such as carbachol (CCh), to the extracellular recording medium (Brombas et al., 2014; Chen et al., 2015; Eggermann and Feldmeyer, 2009; Gulledge and Stuart, 2005; Gulledge et al., 2007; Kawaguchi, 1997; Levy et al., 2008; Poorthuis et al., 2018; Urban-Ciecko et al., 2018; Wang and McCormick, 1993). Emerging data suggest that ACh controls the excitability of neocortical neurons, enhances the signal-to-noise ratio of cortical responses, and modifies the threshold for activity-dependent synaptic modifications by activating postsynaptic muscarinic (mAChR) or nicotinic (nAChR) receptors (Couey et al., 2007; Disney and Aoki, 2008; Herrero et al., 2008; Metherate, 2004; Minces et al., 2017; Poorthuis et al., 2013). At the cellular level, it is understood that ACh mostly activates mAChRs to depolarize neurons and initiate action potentials (APs) (Eggermann et al., 2014; Kawaguchi, 1997; Krnjević et al., 1971). However, a handful of studies also suggest that ACh transiently activates mAChRs and strongly inhibits the initiation of APs in neocortical pyramidal neurons (Gulledge and Stuart, 2005; Gulledge et al., 2007). At the level of synapses, it is known that ACh reduces the efficacy of excitatory connections in the neocortex. For example, in synaptic connections between thick-tufted layer 5 pyramidal cells (TTPCs), which are marked with pronounced short-term depression, bath-application of 5–10 μM of CCh following presynaptic stimulation, rapidly reduces the rate of depression in a train of postsynaptic potentials (PSPs) without affecting the so-called stationary PSPs (Levy et al., 2008; Tsodyks and Markram, 1997). In contrast, administering a similar amount of CCh on facilitating synaptic connections between TTPCs and Martinotti cells (MCs) increases the strength of successive PSPs (Levy et al., 2008). Although some of the cell-type, and connection-type specific effects of ACh in the neocortex have been experimentally mapped, the vast majority remains unknown.

It is thought that ACh is crucially implicated in cognitive functions, ranging from arousal, sleep-wake cycles, learning, and memory among others, yet it has been difficult to develop a unifying view of how ACh controls neuronal and synaptic physiology and impacts neocortical network dynamics. An impediment in this direction is probably due to the fact that ACh differentially controls the activity of neocortical neurons and synapses in complex ways, making it difficult to reconcile its systemic effects (Muñoz and Rudy, 2014). Computational models of neocortical microcircuitry at the cellular and synaptic level of biological detail not only offer an integrative platform to bring together experimental data capturing the specific effects of ACh on dendrites, neurons and synapses, but also make it possible to generate predictions on the actions of ACh at the network level.

As a way forward, we developed a first-draft, data-driven framework that leverages a recent, rigorously validated digital model of the microcircuitry of juvenile rodent somatosensory cortex ((Markram et al., 2015; Ramaswamy et al., 2015) ; Figure 1) comprising ~31,000 neurons distributed across 6 layers, 55 layer-specific morphological (m), 11 electrical (e) and 207 morpho-electrical (me) neuron subtypes that are connected through ~40 million synapses and 6 dynamical synapse (s) types (Figure 2). Next, we augmented the model by integrating the anatomy of the laminar distribution of cholinergic varicosities (Figure 3), and the phenomenological cell-type specific effects of ACh neuronal and synaptic physiology from published literature (Chen et al., 2015; Eggermann and Feldmeyer, 2009; Gulledge and Stuart, 2005; Gulledge et al., 2007; Kawaguchi, 1997; Levy et al., 2008; Tsodyks and Markram, 1997). This data-driven approach enabled us to bridge how the local impact of ACh on neurons and synapses are broadcast to the global level and influence the emergence of neocortical network states. Using this framework, we derive preliminary predictions, which suggest that a dose-dependent change in ACh levels shifts neocortical network state from highly synchronous to asynchronous activity, and distinctly shapes the structure of spike-spike cross-correlations between specific neuronal populations.

**Figure 1.**
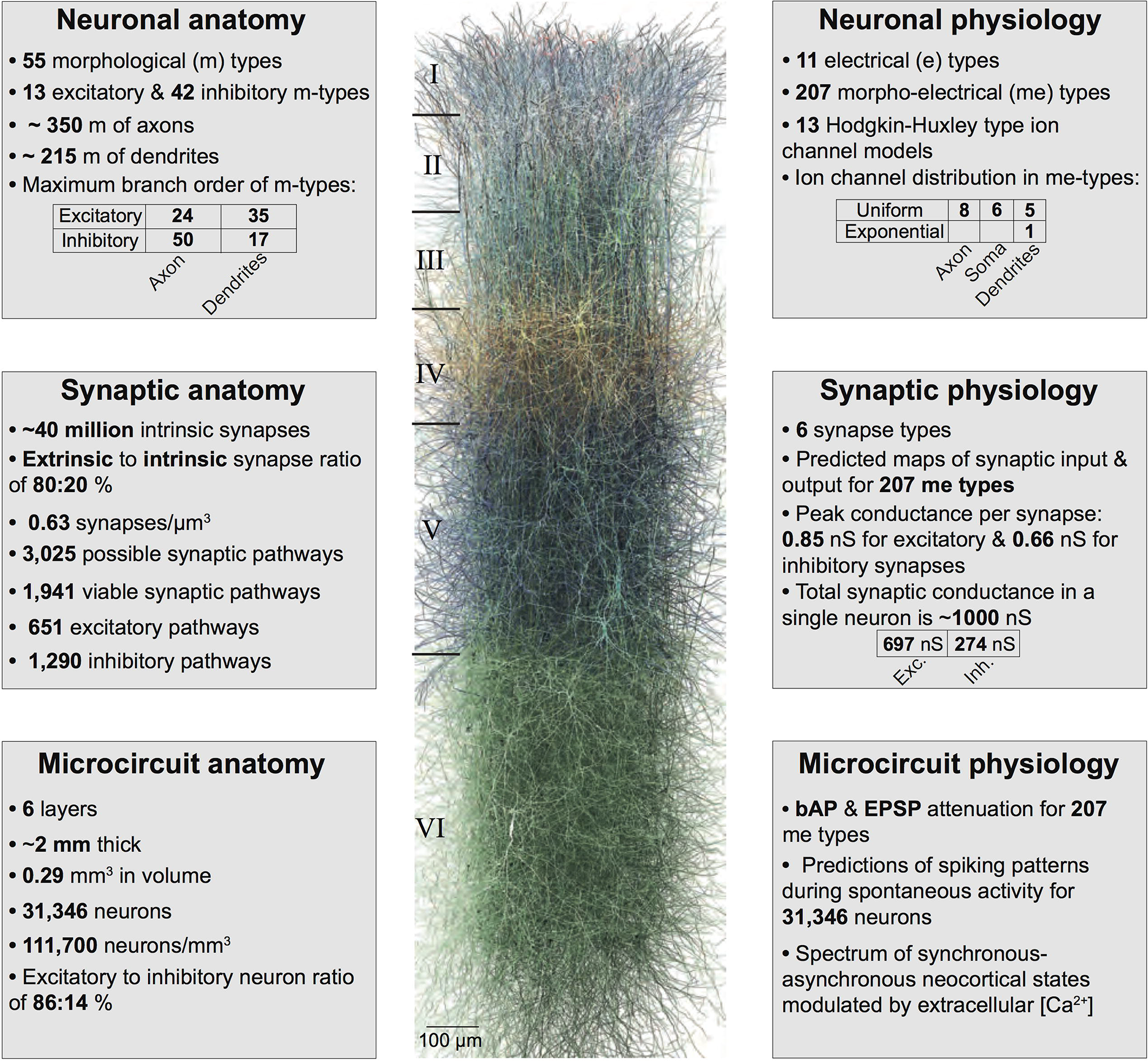
Summary of the biologically detailed tissue model of neocortical microcircuitry. **Top left:** overview of neuronal anatomy in the reconstruction. **Top right:** summary of neuronal physiology. **Middle left:** overview of synaptic anatomy. **Middle right:** fact and figures on synaptic physiology. **Bottom left:** summary of microcircuit anatomy. **Bottom right:** overview of microcircuit physiology.

**Figure 2.**
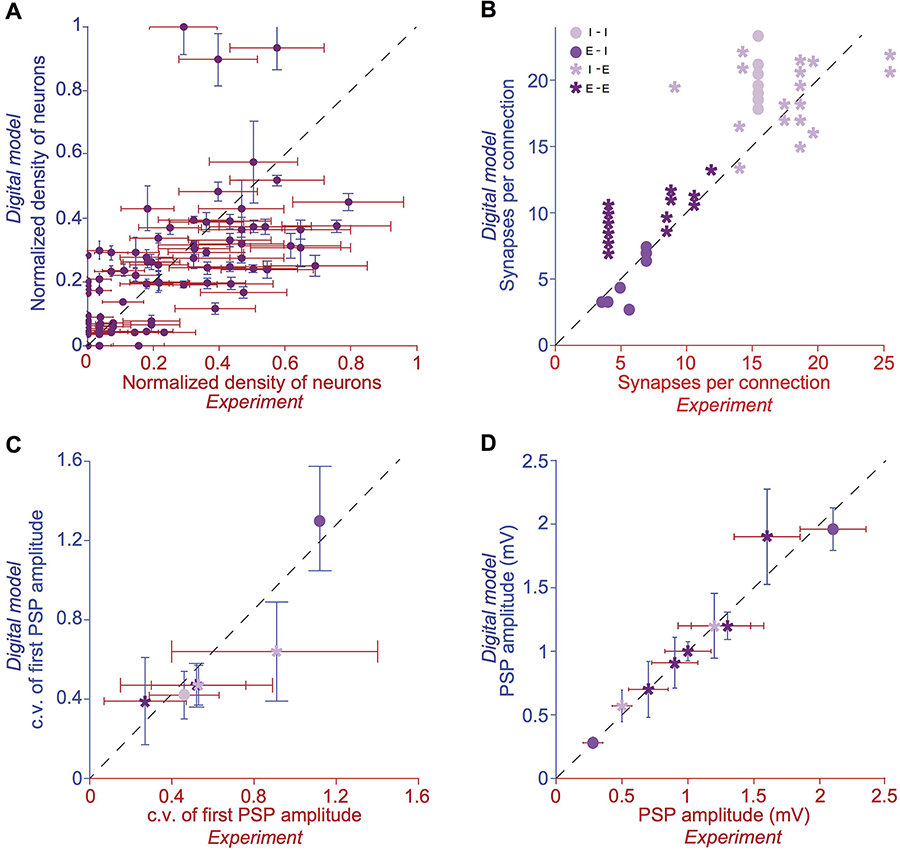
Validation of anatomical and physiological properties in the tissue model of neocortical microcircuitry. **(A)** Normalized neuronal densities. Number of stained neurons per 100 μm bin from layers 1 to 6. Red: *experiment* (counts/bin), blue: *digital model* (counts/bin; mean±SD, N=100 bins). Dashed line has unit slope. **(B)** Mean number of synapses per connection in excitatory-excitatory (E-E), excitatory-inhibitory (E-I), inhibitory-excitatory (I-E), and inhibitory-inhibitory (I-I) pathways. Red: experiment, blue: digital model. Dashed line has unit slope. **(C)** mean coefficient of variation (c.v.; defined as standard deviation/mean) of the amplitude of the postsynaptic potential for pathways some of the pathways in (B). **(D)** same as in C, but for the mean amplitude of the postsynaptic potential for some of the pathways in (B).

**Figure 3.**
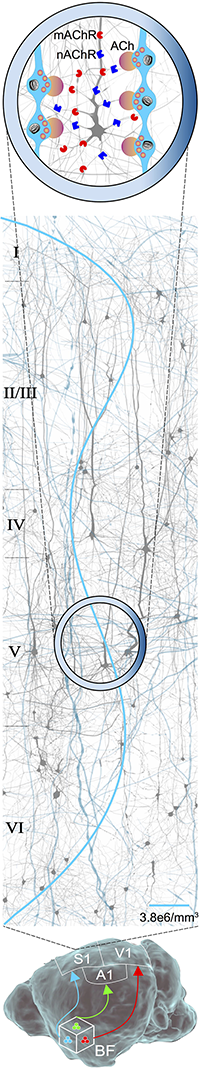
Schematic of cholinergic innervation of neocortical microcircuitry. **Bottom:** Illustration of cholinergic projections from the basal forebrain (BF) to the primary sensory cortices: somatosensory (S1), visual (V1), and auditory (A1). **Middle:** schematic of a prototypical neocortical microcircuit. A forest of cholinergic axons innervating the microcircuit (in cyan) is shown in the background. The average laminar profile of varicosities formed by an ascending cholinergic axon (cyan line) is depicted. **Top:** zoomed-in view of layer 5 with a cartoon of cholinergic varicosities (cyan) releasing ACh (in orange/purple). Muscarinic (mAChR; in red) and nicotinic (nAChR; in blue) receptors are distributed on the dendrites of an exemplar L5 TTPC in the background.

## Results

### Cholinergic innervation of neocortical microcircuitry

The anatomy of long-range axons of cholinergic neurons projecting to the neocortex controls ACh release, which regulates network activity. Therefore, as the first step, we began by integrating anatomical data on the density distribution of cholinergic varicosities across all layers in developing rat somatosensory cortex (Mechawar et al., 2000). The mean density of cholinergic varicosities was the highest in layer 1 (5*10^6^/mm^3^), and the lowest in layer 4 (2.9*10^6^/mm^3^). The overall density of cholinergic varicosities across all neocortical layers was 3.8*10^6^/mm^3^. Based on the innervation profile of cholinergic varicosities across all layers within a prototypical neocortical microcircuit with a volume of ~0.3 mm^3^ (Figure 3), and previous estimates of an average cholinergic axon length of 10 m/mm^3^ (Gritti et al., 1993; Mechawar et al., 2000; Rye et al., 1984), we predict that the entire rat primary somatosensory cortex with an approximate volume of 35 mm^3^, is innervated by ~0.4 km of cholinergic axons containing about 162 million varicosities. It is estimated that about 1000–2000 cholinergic neurons project to the somatosensory cortex from the NBM (Gritti et al., 1993; Rye et al., 1984). Therefore, it could be inferred that the axonal arbor of each cholinergic neuron averages about 0.5–1 m in length and accommodates about 100,000–200,000 varicosities.

### ACh modulation of neuronal physiology

As the next step, we integrated experimental data on the impact of ACh on the resting membrane potential and cellular excitability of neocortical neurons, which enabled us to build a dose-dependent profile across a range of ACh concentrations (Figure 4A, B; (Chen et al., 2015; Eggermann and Feldmeyer, 2009; Gulledge and Stuart, 2005; Gulledge et al., 2007 Levy et al., 2008)). We have previously shown that a piece of neocortical tissue, ~0.3 mm^3^ in volume, consists of 55 m-types and 11 e-types, resulting in 207 me-types distributed across 6 layers (Figure 1). Next, we used validated digital models of 207 me-types that were optimized to reproduce diverse electrophysiological features of excitatory and inhibitory neocortical neurons such as AP amplitudes and widths, mean firing frequency and accommodation index (Ramaswamy et al., 2015; Van Geit et al., 2016). We extended these models by identifying an appropriate level of depolarizing somatic step current injection, which led to an increase in the resting membrane potential and firing frequency of each me-type to mimic the dose-dependent effects of ACh on cellular excitability (Figure 4D; see Methods). The amount of injected step current used to simulate cellular excitability at different ACh levels was expressed in terms of percentage of the minimum current injection required for each me-type model to generate at least a single AP (rheobase; see Methods). In this first-draft implementation, we assumed that all me-types showed the same dose-dependent effects. All me-types responded with a change in intrinsic excitability that was predicted to switch from sub-threshold to supra-threshold behavior at an ACh concentration of ~50 μM (Figure 4D; six randomly chosen me-types are shown). The mean AP firing frequency in all me-types increased significantly from 5 to 10 Hz for a four-fold change in ACh from 50 to 200 μM (Figure 4D).

**Figure 4.**
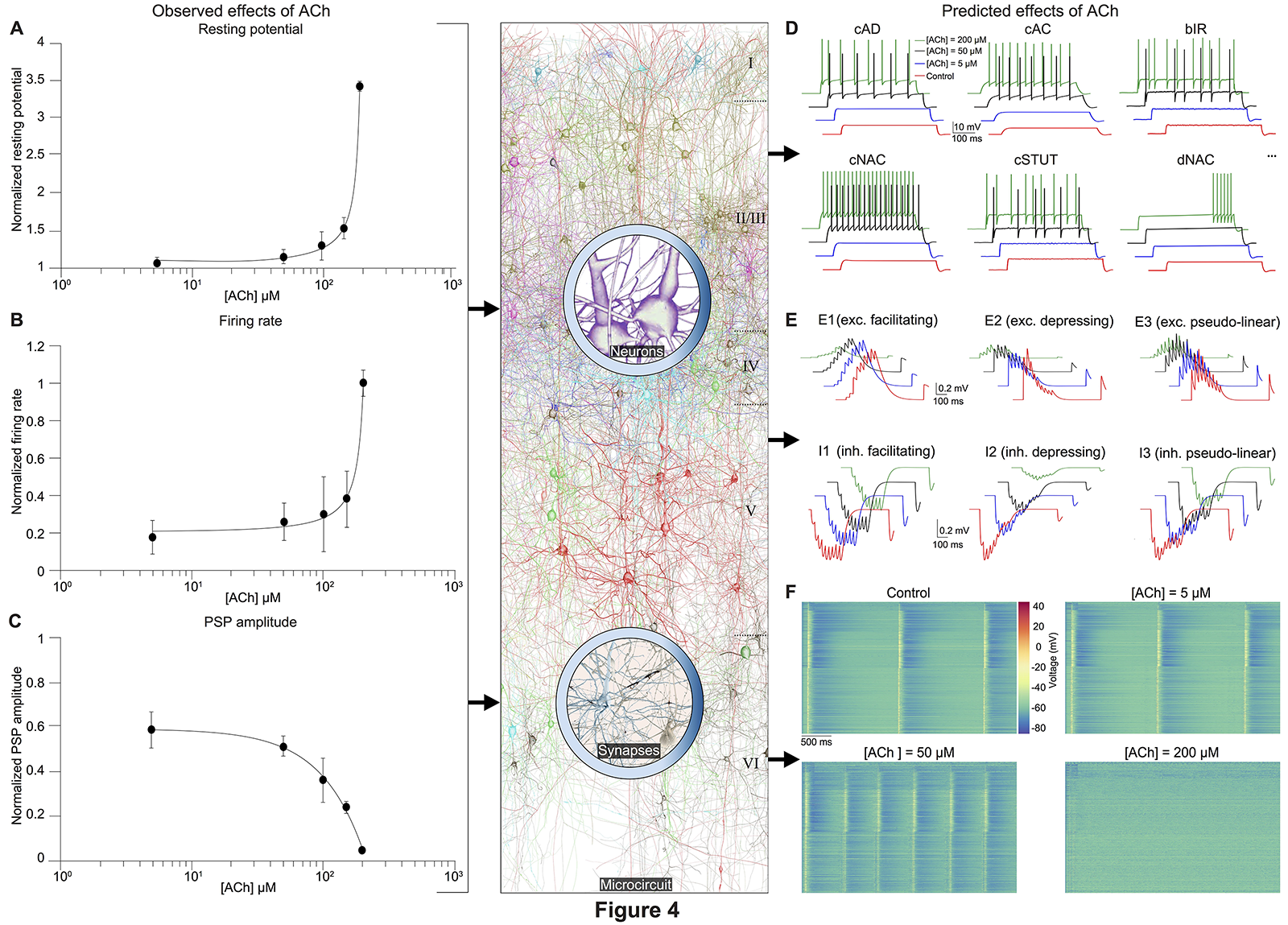
Integrated summary of the cellular, synaptic and microcircuit effects of ACh in the tissue model of neocortical microcircuitry. **(A)** Integrated sparse data-sets from published literature on the dose-response effects of ACh on the resting membrane potential of neocortical neurons (normalized). **(B)** same as in A, but for neuronal firing rate. **(C)** same as in A, but for the amplitude of postsynaptic potentials. sensitivity of neuronal and synaptic physiology. **(D)** Predicted effects of different ACh levels on the resting potential and firing rate of neocortical e-types. Only six e-types are shown. cAD - continuous accommodating (pyramidal), cAC - continuous accommodating (interneurons), bIR - burst irregular, cNAC - continuous non-accommodating, cSTUT - continuous stuttering, dNAC - delayed non-accommodating. **(E)** Predicted effects of different ACh levels on the physiology of all neocortical s-types. **(F)** Prediction of the effect of ACh concentration on network dynamics. Clockwise from left, voltage rasters of 1,000 randomly sampled neurons across layers 1–6 at different ACh concentrations.

### ACh control of synaptic physiology

As the final step, we unified relevant published data, and extrapolated a dose-dependent activation curve of the effects of varying concentrations of ACh on the response amplitude of the first PSP for all neocortical s-types (Figure 4C). It is known that neocortical synapses exhibit at least six distinct forms of excitatory (E) and inhibitory (I) short-term dynamics that are used to distinguish synaptic connections into facilitating (E1 and I1), depressing (E2 and I2), and pseudo-linear (E3 and I3) s-types (refs). We have previously shown that 55 m-types establish around 1,941 morphology-specific synaptic connection types, whose short-term dynamics are governed by one of the 6 s-types dictated by the prepost combination of m-types (Markram et al., 2015; Ramaswamy et al., 2015; Reimann et al., 2015). We augmented this model to include the effects of ACh modulation of the PSP amplitude of s-types and derived predictions on how their short-term facilitating, depressing, and pseudo-linear dynamics are controlled by ACh (Figure 4E). It is known that ACh powerfully modulates the PSP amplitude of synaptic connections between excitatory neocortical m-types, very likely by modifying the probability of glutamate release (Eggermann and Feldmeyer, 2009; Levy et al., 2008). However, it remains unclear if ACh controls inhibitory synaptic transmission in the neocortex by modulating GABA release in similar ways to glutamate (Kruglikov and Rudy, 2008; Yamamoto et al., 2010). Therefore, in this first-draft implementation, we assumed that ACh regulates the release of both glutamate and GABA in comparable ways (Figure 4C).

In order to simulate the change in PSP amplitude as a function of ACh concentration, we modified the neurotransmitter release probability for all synaptic contacts underlying m-type specific connections according the extrapolated dose-dependence curve compiled from literature (Figure 4C). We found that ACh exerted highly diverse effects on the PSP amplitude for the 6 s-types (Figure 4E). The impact of ACh concentrations (5–200 μM) on the first PSP amplitude evoked by a train of 9 presynaptic APs at 30 Hz was superficial compared to control for both E1 (between a L2/3 PC and a MC) and I1 (between a L2/3 small basket cell (SBC) and a PC) s-types (Figure 4E, top left; maximum responses are normalized to control). However, the very pronounced facilitation typically observed for the E1 s-type was strongly suppressed at higher (200 μM), rather than lower concentrations (5–100 μM) of ACh (Figure 4E, top left). The amplitude of the first PSP and the subsequent facilitating dynamics for the I1 s-type was not substantially modulated by ACh, albeit a four-fold increase in concentration (Figure 4E, bottom left; from 5 μM to 200 μM). The physiology of both E2 (between L5 two thick-tufted pyramidal cells; Figure 4E, top center) and I2 (between a L5 MC and a TTPC; Figure 4E, bottom center) s-types was crucially impacted by different ACh levels (5–200 μM). On average, the first PSP amplitude for both E2 and I2 s-types was reduced to about 75%, 50%, and 10% of control at ACh concentrations of 5, 50 and 200 μM ACh, respectively (Figure 4E, top and bottom center). Amplitude of subsequent PSPs decreased to 50–80% of control with markedly diminished rates of depression, but consistent with previous observations, did not critically impact the amplitude of stationary PSPs (Eggermann and Feldmeyer, 2009; Levy et al., 2008; Tsodyks and Markram, 1997). Higher concentrations of ACh at 200 μM almost completely shutoff depressing synaptic transmission (Figure 4E, top and bottom center). For E3 (between two L6 PCs; Figure 4E, top right) and I3 (between a L5 Nest basket cell (NBC) and a TTPC; Figure 4E, bottom right) pseudo-linear s-types ACh concentrations between 5–100 μM did not cause a conspicuous increase in the amplitude of the first PSP in a train. At ACh concentrations of 5 and 50 μM, the mean amplitude of the first PSP for E3 and I3 s-types was approximately 70% and 85% of control, respectively (Figure 4E, top and bottom right).

Whereas, the amplitude of the first PSP at 200 μM ACh was powerfully modulated and diminished to about 10% and 50% for E3 and I3 s-types, respectively (Figure 4E, top and bottom right). However, despite an exponential increase in ACh levels from 5 μM to 200 μM the modulation of pseudo-linear dynamics for E3 and I3 s-types appeared to be insensitive to ACh. We predict that an increase in ACh concentration, more than an order of magnitude, has a steep modulatory effect on the physiology of E2 and I2 s-types, but only a superficial impact on E1, I1, E3, and I3 s-types.

### ACh modulation of network states

It is thought that ACh enhances arousal and vigilance in primary sensory cortices by altering the signal-to-noise ratio of incoming synaptic input (Minces et al., 2017). However, it remains unclear how the differential regulation of neuronal and synaptic physiology by ACh, specifically the modulation of feedforward excitatory and feedback inhibitory transmission, influences the emergence of neocortical network states. In order to explore how the local effects of ACh on cells and synapses modulate global network states, we incorporated phenomenological models of ACh control of neuronal and synaptic physiology into a validated biologically detailed reconstruction of neocortical microcircuitry and simulated the effects of dose-dependent ACh modulation on network activity (ref). To enable a direct comparison with experimental data obtained from cortical slices on the impact of ACh on cellular excitability and synaptic transmission, we created a virtual slice (with a thickness of ~231 μm; see Methods) to explore neocortical network states for a range of ACh concentrations (see Methods). We simulated spontaneous activity in the virtual slice by applying tonic background depolarization and found that in the control condition without any extracellular ACh, neocortical network activity exhibited low-frequency (~1.7 Hz), highly synchronous bursts of oscillatory behavior (Figure 4F, top left) akin to slow-wave sleep (Sanchez-Vives and McCormick, 2000; Steriade et al., 1993). ACh concentrations at 5 μM and 50 μM further diminished the frequency of synchronous oscillatory network activity (Figure 4F, top right and bottom right). At ACh levels of 200 μM, slow oscillatory bursts of synchronous network activity were superseded by irregular asynchronous activity, resembling active waking states (Figure 4F, bottom right). The transition from synchronous to asynchronous neocortical states occurred at ~75 μM. Interestingly, we found that a change in < 50 μM of ACh can switch neocortical dynamics from the synchronous to asynchronous state, divulging two distinct network activity regimes.

Next, we investigated the effect of ACh concentrations in shaping spike-time cross-correlations for pairs of neurons - excitatory-excitatory (L23 PC-L23 PC; Figure 5A), excitatory-inhibitory (L4 PC-L4 NBC; Figure 5B), inhibitory-excitatory (L5 NBC-L5 TTPC; Figure 5C), and inhibitory-inhibitory (L6 MC-L6 MC; Figure 5D). We observed a striking diversity in the average cross-correlation profiles for different pairs of neurons comprising these populations, which was computed as the mean spike-time cross-correlation from 10,000–20,000 randomly sampled pairs. At the outset, correlations differed in their temporal profiles (Figure 5A-D). Upon closer examination of these correlation profiles, in particular with the peak lag (delay to peak) and the median lag (delay of the median) revealed that they differed significantly between all examined populations (Figure 5A–D). For example, between pairs of excitatory neurons, the cross-correlations at different ACh concentrations were similar to auto-correlations, with a very small range in peak lag values (Figure 5A) in comparison to the crosscorrelations between excitatory and inhibitory (Figure 5B).

**Figure 5.**
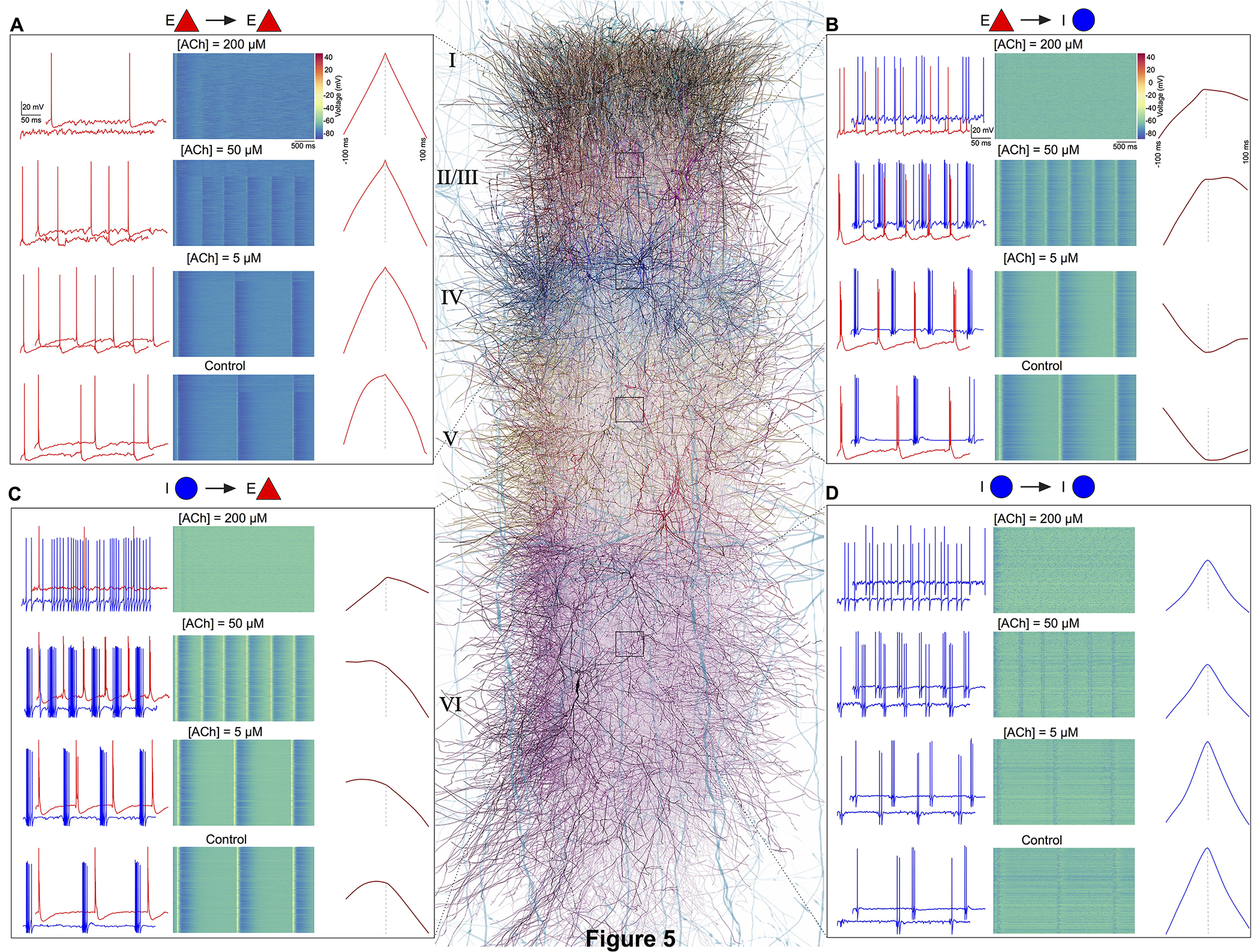
ACh shapes spike-spike cross-correlations. (**A**) **Left:** spontaneous spiking activity in a randomly chosen pair of excitatory-excitatory (L23 PC-L23 PC) neurons at different ACh levels. **Middle:** voltage raster for all L23 PCs at different ACh levels. **Right:** spike-time cross correlations for 10,000–20,000 randomly sampled pairs of L23PCs at different ACh levels. **(B)** same as in A, but for excitatory-inhibitory neurons (L4 PC-L4 NBC). **(C)** same as in A, but for inhibitory-excitatory neurons (L5 NBC-L5 TTPC). **(D)** same as in A, but for inhibitory-inhibitory neurons (L6 MC-L6 MC).

We, therefore, predict that endogenous ACh released from subcortical structures, such as the NBM, powerfully modulates neocortical activity giving rise to a spectrum of network states ranging from one extreme where low ACh levels bring about synchronous activity, to another, where high ACh concentrations lead to asynchrony. Our results are broadly consistent with studies employing optogenetic approaches that associate sleep states with low ACh levels and wakefulness with high ACh concentrations (Chen et al., 2015; Lee and Dan, 2012). We have previously demonstrated that extracellular calcium (Ca^2+^) regulates the emergence of synchronous and asynchronous network states in the neocortex (Markram et al., 2015). Based on these preliminary results, we hypothesize that neuromodulators, such as ACh, provide a complementary functional mechanism in the neocortex, similar to a “push-pull” switch, where the interplay of low ACh and high Ca^2+^ pushes network state towards synchronous activity, whereas high ACh and low Ca^2+^ levels pulls network activity towards asynchrony. We propose that ACh orchestrates neocortical dynamics by generating a spectrum of network states - where a regime of correlated firing in neurons causes synchronous activity that could modulate functions such as coincidence detection, response selection and binding, and asynchronous activity could promote encoding of new information boosted by heightened attention to incoming sensory input.

## Discussion

This study presents a first-draft implementation of a data-driven framework, which unifies the phenomenological effects of the regulation of local cellular excitability and synaptic physiology by neuromodulators to predict their global impact on the emergence of neocortical network states. As a first foray into exploring the utility of this framework, we integrated biological data on how ACh controls the electrical and synaptic properties of cell-types in the rodent neocortex and derived preliminary insights into how a range of ACh levels generated a spectrum of network states.

Emerging experimental state-of-the-art suggests that ACh exerts a divergent control of neocortical neurons and synapses. The effects of ACh on the vast majority of neurons and synapses in the neocortex remains unknown. It is unlikely that the myriad forms by which ACh could control the physiology of neurons and synapses will be unambiguously understood in the future. However, the advent of optogenetics to interrogate the cell-type specific effects of ACh combined with a data-driven computational framework, such as the one presented here holds promise in filling knowledge gaps and accelerating our understanding of the complex spatiotemporal actions of ACh in the neocortex.

Albeit this framework can already provide elementary insights into ACh regulation of neocortical states by bridging cellular, synaptic and network levels, it is still a first-draft and lacks numerous biological details on the anatomy and physiology of ACh innervation of neocortical layers and neurons. Indeed, in this first-draft implementation, we assumed that the dose-dependent activation profile of ACh is homogeneous on excitatory and inhibitory cell-types and their synaptic connections, which is a gross generalization. As the next step to refine the biological accuracy and specificity of our framework, we plan to systematically incorporate laminar data on cholinergic axon lengths, bouton densities, receptor localization and kinetics of ACh receptors, and their impact on the neuronal and synaptic physiology of an assortment of neocortical cell-types.

A methodical integration of biological data on neuromodulatory control of neocortical dynamics into the unifying framework will enable the identification of the unknowns, reconciliation of disparate datasets, and prediction of their general organizing principles. Additionally, our framework not only allows further investigation on the role of ACh in regulating neocortical dynamics but can also be applicable to hypothesize and predict the function of other major neuromodulators - noradrenaline, dopamine, serotonin, and histamine - that influence the emergence of network states. In conclusion, we propose the framework as a complementary resource to existing experimental approaches to advance our understanding of how neuromodulatory systems differentially regulate the activity of a diversity of neurons and synapses and sculpt neocortical network states.

## Methods

A digital model of the microcircuitry of juvenile rodent somatosensory cortex was reconstructed as previously described (Markram et al., 2015; Ramaswamy et al., 2015; Reimann et al., 2015). In brief, the reconstruction process comprised the following:

**Microcircuit dimensions:** Thicknesses of individual layers and the diameter of the microcircuit were used to construct a virtual hexagonal prism. A virtual slice was generated from a 1×7 mosaic of microcircuits as a cortical sheet with a thickness of 230.9 μm and a width of 2800 μm.

**Cellular composition:** Measurements of neuronal densities across neocortical layers and fractions of m- and me-types were used to generate the position of individual neurons in the reconstructed microcircuit, constrained by layer-specific proportions of excitatory and inhibitory neurons. Each neuron was assigned the optimal morphology for its location in the microcircuit.

**Digital neuron morphologies:** Neuronal morphologies were obtained from digital 3D reconstructions of biocytin-stained neurons after whole-cell patch-clamp recordings in 300μm-thick, sagittal neocortical slices from juvenile rat hind-limb somatosensory cortex. Severed neurites of morphologies due to the slicing procedure were algorithmically regrown (Anwar et al., 2009). Neurites were digitally unraveled to compensate for shrinkage. Neuronal morphologies were then cloned to obtain a sufficient representation of all m-types.

**Electrical Neuron Models:** Conductance based, multi-compartmental electrical models of neurons were produced using up to 13 active ion channel mechanisms and a model of intracellular Ca^2+^ dynamics. Axon initial segments (AIS), somata, basal and apical dendrites were modeled as separate, but interconnected compartments. Pyramidal neurons contained two dendritic regions, whereas interneurons contained only one dendritic region. Each region received a separate set of ion channels ((see NMC portal; https://bbp.epfl.ch/nmc-portal; (Ramaswamy et al., 2015)). Only the AIS was simulated. Each AIS was represented by two fixed length sections, each with a length of 30 μm; diameters were obtained from the reconstructed morphology used for model fitting. APs detected in the AIS were propagated to the synaptic contacts with a delay corresponding to the axonal delay required to propagate to each synapse, assuming an axonal velocity of 300 μm/ms. As previously described, electrical models were fitted using a feature-based multi-objective optimization method.

**Synaptic anatomy:** The number and location of synaptic contacts were derived using an algorithm, described previously (Reimann et al., 2015). The algorithm removes axo-dendritic appositions that do not obey the multi-synapse and plasticity reserve rules and ensures compatibility with biological bouton densities.

**Synaptic physiology:** Excitatory synaptic transmission was modeled using both AMPA and NMDA receptor kinetics. Inhibitory synaptic transmission was modeled with a combination of GABA_A_ and GABA_B_ receptor kinetics. Stochastic synaptic transmission was implemented as a two-state Markov model of neurotransmitter release, a stochastic implementation of the Tsodyks-Markram dynamic synapse model. Biological parameter ranges for the three model parameters - neurotransmitter release probability, recovery from depression and facilitation - were obtained from experimental measurement for synaptic connections between specific m- and me-types or between larger categories of pre and postsynaptic neurons. Spontaneous miniature PSCs were modeled by implementing an independent Poisson process for each individual synapse that triggered release at rates (λ_spont_) determined by the experimental data.

**Microcircuit simulation:** The digital model of neocortical microcircuitry was simulated using the NEURON simulation enviroment, augmented for execution on a supercomputer (Hines and Carnevale, 1997; Hines et al., 2008a, 2008b), along with custom tools to setup and configure microcircuit simulations, and read output results.

**Implementation of cholinergic afferents:** We used published data on the density of cholinergic varicosities measured in the parietal lobe, which also includes the somatosensory cortex, to generate a laminar density profile for extrinsic cholinergic input (Mechawar et al., 2000). The implementation followed a previously described method (Markram et al., 2015). First, the total thickness of all neocortical layers, which is a measure of vertical depth, was digitized and binned with a bin size of 50 μm. For each depth bin, we then found all morphological segments contained inside that bin and then continuously drew random segments from the pool and generated a varicosity at their centers.

**Implementation of dose-dependent effects of ACh on cellular excitability:** Dose-dependent effects of ACh on cellular excitability was achieved by depolarizing somatic step current injection, which caused an increase in the resting membrane potential and firing frequency. Step currents were expressed in terms of percentage of the minimum step current injection required for each cell to spike at least once (rheobase).

**Implementation of dose-dependent effects of ACh on synaptic transmission:** Dose-dependent effects of ACh on synaptic physiology was achieved by changing the utilization of synaptic efficacy parameter (U) in the stochastic synapse model. The effect of ACh on excitatory and inhibitory synaptic response amplitudes were simulated by modifying the neurotransmitter release probability for all synaptic contacts underlying m-type specific connections according the extrapolated dose-dependence curve compiled from literature (Figure 4C). Due to lack of data for specific synaptic connection-types, we assumed that all excitatory and inhibitory connections showed the same dose-dependent effects to ACh.

**Cross-correlations:** Mean spike-spike cross-correlations were computed as the average of all spike-times measured in 10,000–20,000 randomly sampled pairs of excitatory-excitatory, excitatory-inhibitory, inhibitory-excitatory, and inhibitory-inhibitory neurons. Cross-correlograms (Figure 5A-D) were computed in Matlab (version 9.1).

## Acknowledgements

This work was supported by funding from the ETH Domain for the Blue Brain Project (BBP); The Human Brain Project through the European Union Seventh Framework Program (FP7/2007–2013) under grant agreement no. 604102 (HBP) and from the European Union H2020 FET program through grant agreement no. 720270 (HBP SGA1); The Cajal Blue Brain Project (MINECO); The BlueBrain IV BlueGene/Q system is financed by ETH Board Funding to the Blue Brain Project as a National Research Infrastructure and hosted at the Swiss National Supercomputing Center (CSCS).

## References

Anwar, H., Riachi, I., Hill, S., Schurmann, F., and Markram, H. (2009). An approach to capturing neuron morphological diversity. In Computational Modeling Methods for Neuroscientists, (The MIT Press), pp. 211–231.

Brombas, A., Fletcher, L.N., and Williams, S.R. (2014). Activity-Dependent Modulation of Layer 1 Inhibitory Neocortical Circuits by Acetylcholine. J. Neurosci. 34, 1932–1941.

Chen, N., Sugihara, H., and Sur, M. (2015). An acetylcholine-activated microcircuit drives temporal dynamics of cortical activity. Nat. Neurosci. 18, 892–902.

Couey, J.J., Meredith, R.M., Spijker, S., Poorthuis, R.B., Smit, A.B., Brussaard, A.B., and Mansvelder, H.D. (2007). Distributed network actions by nicotine increase the threshold for spike-timing-dependent plasticity in prefrontal cortex. Neuron 54, 73–87.

Disney, A.A., and Aoki, C. (2008). Muscarinic acetylcholine receptors in macaque V1 are most frequently expressed by parvalbumin-immunoreactive neurons. J. Comp. Neurol. 507, 1748–1762.

Eggermann, E., and Feldmeyer, D. (2009). Cholinergic filtering in the recurrent excitatory microcircuit of cortical layer 4. Proc. Natl. Acad. Sci. 106, 11753–11758.

Eggermann, E., Kremer, Y., Crochet, S., and Petersen, C.C.H. (2014). Cholinergic Signals in Mouse Barrel Cortex during Active Whisker Sensing. Cell Rep. 9, 1654–1660.

Gielow, M.R., and Zaborszky, L. (2017). The Input-Output Relationship of the Cholinergic Basal Forebrain. Cell Rep. 18, 1817–1830.

Gritti, I., Mainville, L., and Jones, B.E. (1993). Codistribution of GABA- with acetylcholine-synthesizing neurons in the basal forebrain of the rat. J. Comp. Neurol. 329, 438–457.

Gulledge, A.T., and Stuart, G.J. (2005). Cholinergic inhibition of neocortical pyramidal neurons. J. Neurosci. Off. J. Soc. Neurosci. 25, 10308–10320.

Gulledge, A.T., Park, S.B., Kawaguchi, Y., and Stuart, G.J. (2007). Heterogeneity of phasic cholinergic signaling in neocortical neurons. J. Neurophysiol. 97, 2215–2229.

Hasselmo, M.E. (1995). Neuromodulation and cortical function: modeling the physiological basis of behavior. Behav. Brain Res. 67, 1–27.

Hasselmo, M.E. (1999). Neuromodulation: acetylcholine and memory consolidation. Trends Cogn. Sci. 3, 351–359.

Herrero, J.L., Roberts, M.J., Delicato, L.S., Gieselmann, M.A., Dayan, P., and Thiele, A. (2008). Acetylcholine contributes through muscarinic receptors to attentional modulation in V1. Nature 454, 1110–1114.

Hines, M.L., and Carnevale, N.T. (1997). The NEURON Simulation Environment. Neural Comput. 9, 1179–1209.

Hines, M.L., Markram, H., and Schurmann, F. (2008a). Fully implicit parallel simulation of single neurons. J. Comput. Neurosci. 25, 439–448.

Hines, M.L., Eichner, H., and Schurmann, F. (2008b). Neuron splitting in compute-bound parallel network simulations enables runtime scaling with twice as many processors. J. Comput. Neurosci. 25, 203–210.

Juliano, S.L., and Jacobs, S.E. (1995). The Role of Acetylcholine in Barrel Cortex. In The Barrel Cortex of Rodents, (Springer, Boston, MA), pp. 411–434.

Kawaguchi, Y. (1997). Selective Cholinergic Modulation of Cortical GABAergic Cell Subtypes. J. Neurophysiol. 78, 1743–1747.

Krnjević, K., Pumain, R., and Renaud, L. (1971). The mechanism of excitation by acetylcholine in the cerebral cortex. J. Physiol. 215, 247–268.

Kruglikov, I., and Rudy, B. (2008). Perisomatic GABA Release and Thalamocortical Integration onto Neocortical Excitatory Cells Are Regulated by Neuromodulators. Neuron 58, 911–924.

Lee, S.-H., and Dan, Y. (2012). Neuromodulation of Brain States. Neuron 76, 209–222.

Levey, A.I., Hallanger, A.E., and Wainer, B.H. (1987). Cholinergic nucleus basalis neurons may influence the cortex via the thalamus. Neurosci. Lett. 74, 7–13.

Levy, R.B., Reyes, A.D., and Aoki, C. (2008). Cholinergic modulation of local pyramid-interneuron synapses exhibiting divergent short-term dynamics in rat sensory cortex. Brain Res. 1215, 97–104.

Markram, H., Muller, E., Ramaswamy, S., Reimann, M.W., Abdellah, M., Aguado Sanchez, C., Ailamaki, A., Alonso-Nanclares, L., Antille, N., Arsever, S., et al. (2015). Reconstruction and Simulation of Neocortical Microcircuitry. Cell 163, 456–492.

McCormick, D.A. (1992). Neurotransmitter actions in the thalamus and cerebral cortex. J. Clin. Neurophysiol. Off. Publ. Am. Electroencephalogr. Soc. 9, 212–223.

Mechawar, N., Cozzari, C., and Descarries, L. (2000). Cholinergic innervation in adult rat cerebral cortex: A quantitative immunocytochemical description. J. Comp. Neurol. 428, 305–318.

Mesulam, M.-M., Mufson, E.J., Levey, A.I., and Wainer, B.H. (1983). Cholinergic innervation of cortex by the basal forebrain: Cytochemistry and cortical connections of the septal area, diagonal band nuclei, nucleus basalis (Substantia innominata), and hypothalamus in the rhesus monkey. J. Comp. Neurol. 214, 170–197.

Metherate, R. (2004). Nicotinic Acetylcholine Receptors in Sensory Cortex. Learn. Mem. 11, 50–59.

Metherate, R., Cox, C.L., and Ashe, J.H. (1992). Cellular bases of neocortical activation: modulation of neural oscillations by the nucleus basalis and endogenous acetylcholine. J. Neurosci. 12, 4701–4711.

Minces, V., Pinto, L., Dan, Y., and Chiba, A.A. (2017). Cholinergic shaping of neural correlations. Proc. Natl. Acad. Sci. 201621493.

Muñoz, W., and Rudy, B. (2014). Spatiotemporal specificity in cholinergic control of neocortical function. Curr. Opin. Neurobiol. 26, 149–160.

Picciotto, M.R., Higley, M.J., and Mineur, Y.S. (2012). Acetylcholine as a Neuromodulator: Cholinergic Signaling Shapes Nervous System Function and Behavior. Neuron 76, 116–129.

Poorthuis, R.B., Bloem, B., Schak, B., Wester, J., Kock, D., J, C.P., and Mansvelder, H.D. (2013). Layer-Specific Modulation of the Prefrontal Cortex by Nicotinic Acetylcholine Receptors. Cereb. Cortex 23, 148–161.

Poorthuis, R.B., Muhammad, K., Wang, M., Verhoog, M.B., Junek, S., Wrana, A., Mansvelder, H.D., and Letzkus, J.J. (2018). Rapid Neuromodulation of Layer 1 Interneurons in Human Neocortex. Cell Rep. 23, 951–958.

Ramaswamy, S., Courcol, J.-D., Abdellah, M., Adaszewski, S., Antille, N., Arsever, S., Guy Antoine, A.K., Bilgili, A., Brukau, Y., Chalimourda, A., et al. (2015). The Neocortical Microcircuit Collaboration Portal: A Resource for Rat Somatosensory Cortex. Front. Neural Circuits 9, 44.

Reimann, M.W., King, J.G., Muller, E.B., Ramaswamy, S., and Markram, H. (2015). An algorithm to predict the connectome of neural microcircuits. Front. Comput. Neurosci. 120.

Rye, D.B., Wainer, B.H., Mesulam, M.M., Mufson, E.J., and Saper, C.B. (1984). Cortical projections arising from the basal forebrain: a study of cholinergic and noncholinergic components employing combined retrograde tracing and immunohistochemical localization of choline acetyltransferase. Neuroscience 13, 627–643.

Sanchez-Vives, M.V., and McCormick, D.A. (2000). Cellular and network mechanisms of rhythmic recurrent activity in neocortex. Nat. Neurosci. 3, 1027–1034.

Steriade, M., Amzica, F., and Nunez, A. (1993). Cholinergic and noradrenergic modulation of the slow (approximately 0.3 Hz) oscillation in neocortical cells. J. Neurophysiol. 70, 1385–1400.

Tsodyks, M.V., and Markram, H. (1997). The Neural Code Between Neocortical Pyramidal Neurons Depends on Neurotransmitter Release Probability. Proc. Natl. Acad. Sci. 94, 719–723.

Urban-Ciecko, J., Jouhanneau, J.-S., Myal, S.E., Poulet, J.F.A., and Barth, A.L. (2018). Precisely Timed Nicotinic Activation Drives SST Inhibition in Neocortical Circuits. Neuron 97, 611–625.e5.

Van Geit, W., Gevaert, M., Chindemi, G., Rössert, C., Courcol, J.-D., Muller, E.B., Schürmann, F., Segev, I., and Markram, H. (2016). BluePyOpt: Leveraging Open Source Software and Cloud Infrastructure to Optimise Model Parameters in Neuroscience. Front. Neuroinformatics 10.

Wang, Z., and McCormick, D.A. (1993). Control of firing mode of corticotectal and corticopontine layer V burst-generating neurons by norepinephrine, acetylcholine, and 1S,3R- ACPD. J. Neurosci. 13, 2199–2216.

Xiang, Z., Huguenard, J.R., and Prince, D.A. (1998). Cholinergic Switching Within Neocortical Inhibitory Networks. Science 281, 985–988.

Yamamoto, K., Koyanagi, Y., Koshikawa, N., and Kobayashi, M. (2010). Postsynaptic Cell Type-Dependent Cholinergic Regulation of GABAergic Synaptic Transmission in Rat Insular Cortex. J. Neurophysiol. 104, 1933–1945.

Zagha, E., and McCormick, D.A. (2014). Neural control of brain state. Curr. Opin. Neurobiol. 29, 178–186.

